# Biophysical and physiological causes of coral reef microbialization

**DOI:** 10.1101/495481

**Authors:** Cynthia B. Silveira, Ty N. F. Roach, Helena Villela, Adam Barno, Brandon Reyes, Esther Rubio-Portillo, Tram Le, Spencer Mead, Mark Hatay, Antoni Luque, Linda Wegley-Kelly, Mark Vermeij, Barbara Bailey, Yui Takeshita, Andreas Haas, Forest Rohwer

**Affiliations:** Department of Biology, San Diego State University, San Diego, CA, USA; Viral Information Institute, San Diego State University, San Diego, CA, USA; Department of Microbiology, Rio de Janeiro Federal University, Rio de Janeiro, Brazil; Department of Physiology, Genetics and Microbiology, University of Alicante, Alicante, Spain; Department of Mathematics & Statistics, San Diego State University, San Diego, CA, USA; CARMABI Foundation, Willemstad, Curaçao; Aquatic Microbiology, Institute for Biodiversity and Ecosystem Dynamics, University of Amsterdam, Amsterdam, The Netherlands; Monterey Bay Aquarium Research Institute, Moss Landing, CA, USA; NIOZ Royal Netherlands Institute for Sea Research and Utrecht University, Texel, The Netherlands

**Keywords:** heterotrophic bacteria, biomass, respiration, carbon metabolism

## Abstract

Coral reefs are declining globally as their primary producer communities shift from stony coral to fleshy macroalgae dominance. Previously, we have shown that the rise of fleshy macroalgae produces dissolved organic carbon (DOC) that lead to microbialization and coral death. Here we test the hypothesis that the biophysical cause of bacterial biomass accumulation is a relative decrease in electron acceptors relative to electron donors due to O_2_ loss from macroalgae. We show that 37 % of photosynthetic O_2_ produced by reef fleshy macroalgae is lost in the form of gas through ebullition from algae surfaces. O_2_ loss increases DOC:O_2_ ratios, decoupling the photosynthetically fixed carbon from oxidative potential for respiration. This biogeochemical environment drives heterotrophic microbes to increase cell-specific DOC consumption and cell sizes, accumulating biomass. In contrast, corals do not lose oxygen and support the growth of smaller and fewer bacteria. *In situ* biomass and metagenomic analyses of 87 reefs across the Pacific and Caribbean show that on algae-dominated reefs bacteria accumulate organic carbon through a Warburg-like increase in aerobic glycolytic metabolism. Because of its biophysical basis, microbialization is predicted to occur in other marine ecosystems shifting primary producer assemblages, such as planktonic communities in warming and acidifying conditions.

## Introduction

Anthropogenic pressures are shifting the composition of reef primary producer communities from calcifying to non-calcifying organisms globally (Cinner et al. 2016; Smith et al. 2016). These changes occur as fleshy algae gain competitive advantage over corals through their interaction with heterotrophic microbes (Barott and Rohwer 2012; Jorissen et al. 2016). High Dissolved Organic Carbon (DOC) release rates by fleshy algae stimulate the growth of the so-called super-heterotrophic bacteria (Haas et al. 2011; Nelson et al. 2013; Kelly et al. 2014). The exacerbated heterotrophic growth at coral-algae interaction zones consume large amounts of oxygen, creating hypoxic zones that kill corals (Haas et al. 2013a; Haas et al. 2014b; Roach et al. 2017). At the reef-scale, shifts from coral to algae dominance stimulate microbial heterotrophic metabolism in the water above the reef (Silveira et al. 2015; Haas et al. 2016; Silveira et al. 2017a). This microbial oxygen consumption creates a feedback loop of coral death, algae overgrowth and microbial biomass accumulation, described as reef microbialization (Dinsdale and Rohwer 2011). Understanding coral reef biomass reallocation into heterotrophic bacterial compartments, in detriment of macro-heterotrophs, requires linking the photosynthesis, respiration, and biomass allocation dynamics during reef degradation (McDole et al. 2012; Haas et al. 2016).

Differences in microbial responses to coral and algae dominance stem from the physiology of these primary producers. Calcifying organisms, including scleractinian corals and crustose coralline algae (CCA), invest 50 to 80 % of their daily fixed carbon in respiration to sustain the energetic costs of calcification (Tremblay et al. 2012). Fleshy macroalgae allocate 10 to 30 % to respiration and release up to 60 % of their primary production as dissolved organic carbon (DOC) (Jokiel and Morrissey 1986; Crossland 1987; Cheshire et al. 1996; Peninsula et al. 2007). The ratio between DOC and O_2_ released by fleshy algae in bottle incubations is higher compared to corals (Haas et al. 2011; Haas et al. 2013b). Coral and algae physiology would predict the opposite pattern because (a) fleshy algae allocate a higher proportion of their daily synthesized carbon to biomass compared to corals, sustaining high herbivory pressure (Falkowski et al. 1984; Duarte and Cebrián 1996; Tremblay et al. 2016), and (b) corals allocate higher proportion of their carbon budget to respiration (Hatcher 1988; Houlbrèque and Ferrier-Pagès 2009; Tremblay et al. 2016). Together, these processes are predicted to increase DOC-to-O_2_ ratios in coral exudates compared to algae, but experimental data has shown the opposite to be true (Haas et al. 2011).

Heterotrophic microbes growing on algal exudates have low growth efficiency, defined as the number of cells produced per unit of carbon consumed (Haas et al. 2011; Nelson et al. 2013; Silveira et al. 2015). Microbes growing on algae-dominated reefs shift their glycolytic metabolism from Embden–Meyerhof–Parnas pathway (EMP) towards Pentose Phosphate (PP) and Entner-Doudoroff (ED) pathways (Haas et al. 2016). All three pathways consume glucose in a series of redox reactions that produce pyruvate. The EMP exclusively produces NADH as reductive potential, while PP and ED store part of the reductive potential in NADPH (Klingner et al. 2015; Spaans et al. 2015). These two reduced coenzymes have different fates in the cell: NADH is mainly used for ATP production via oxidative phosphorylation and NADPH is mainly used for biosynthetic processes (Fuhrer and Sauer 2009). Because oxygen is the final electron acceptor in oxidative phosphorylation, these pathways have distinct O_2_ consumption patterns. Comparative thermodynamic, kinetic and genomic analyses of these pathways demonstrate that the preferential use of each reflect fundamental thermodynamic and ecological constrains and predicts an impact on the cell metabolic budget (Flamholz et al. 2013; Stettner and Segrè 2013).

Beyond primary producer physiology, coral reef organic carbon and oxygen dynamics respond to biophysical properties of these organisms and the surrounding environment. Fleshy turf and macroalgae release up to three times more DOC per photosynthetically produced O_2_ than calcifying organisms (Naumann et al. 2010; Haas et al. 2011; Nelson et al. 2013; Silveira et al. 2015). Yet, algae-dominated systems have consistently lower concentrations of DOC and more extreme daily fluctuations in O_2_ concentrations (Dinsdale et al. 2008; Martinez et al. 2012; Nelson et al. 2013). Fleshy algae produce O_2_ bubbles through heterogeneous nucleation resulting from O_2_ super-saturation at the algae’s surface (Kraines et al. 1996). The gaseous O_2_ in bubbles is not detected by dissolved O_2_ instruments, causing underestimation of autotrophy in oxygen-based primary productivity methods (Atkinson and Grigg 1984; Kraines et al. 1996). While several studies recognize bubble formation on algal surfaces, the fraction of photosynthetic oxygen lost as bubbles from different benthic primary producers has never been quantified (Odum and Odum 1955; Freeman et al. 2018).

Here we test the hypothesis that the relative decrease in electron acceptors relative to electron donors due to O_2_ loss through ebullition selects for heterotrophic bacterial metabolisms on degraded coral reefs. To answer this question, we developed novel chamber incubation experiments and showed that fleshy macroalgae release larger amounts of photosynthetic O_2_ by ebullition compared to corals. As O_2_ comes out of solution, it is no longer available for biological respiration in the water column, increasing the ratio of reducing-to-oxidizing equivalents. We then quantified the microbial physiological responses to these conditions, and observed an in increase in cell volume and organic carbon consumption per cell, and a switch to anabolic metabolism in algae incubations. Finally, we show that the patterns identified in our incubations are observed *in situ*, with an increase in bacterial biomass and a shift toward anabolic metabolism across 87 reefs in the Pacific and Caribbean. These metabolic transitions are the mechanistic basis of microbialization that intensifies coral reef decline.

## Methods

### O_2_ release by benthic primary producers

The rates of dissolved and gaseous O_2_ production by benthic primary producers were measured in two tank experiments (POP Experiments 1 and 2). In both experiments, organisms were incubated in custom-made chambers named peripheral oxygen production (POP) bottles (Figure 1). POP bottles are bell-shaped Polyethylene Terephthalate (PET) bottles with a removable base and two sampling ports, one for dissolved analyte sampling, and one at the top for gas sampling. Primary producers were placed at the bottom and bubbles released from their surfaces during incubation accumulated at the top. Primary producers were placed on the base of the POP bottles, inside a larger tank filled with reef water. The bell-shaped container was then placed over the base, and the bottle was sealed. At the end of the incubation, the gas accumulated at the top of the bottles was collected in a syringe and the volume of gas was recorded. The gas was transferred to a wide-mouth container and O_2_ partial pressure was measured using a polarographic probe (Extech 407510) immediately upon collection. Dissolved O_2_ was determined using a Hatch-Lange HQ40 DO probe.

**Figure 1.**
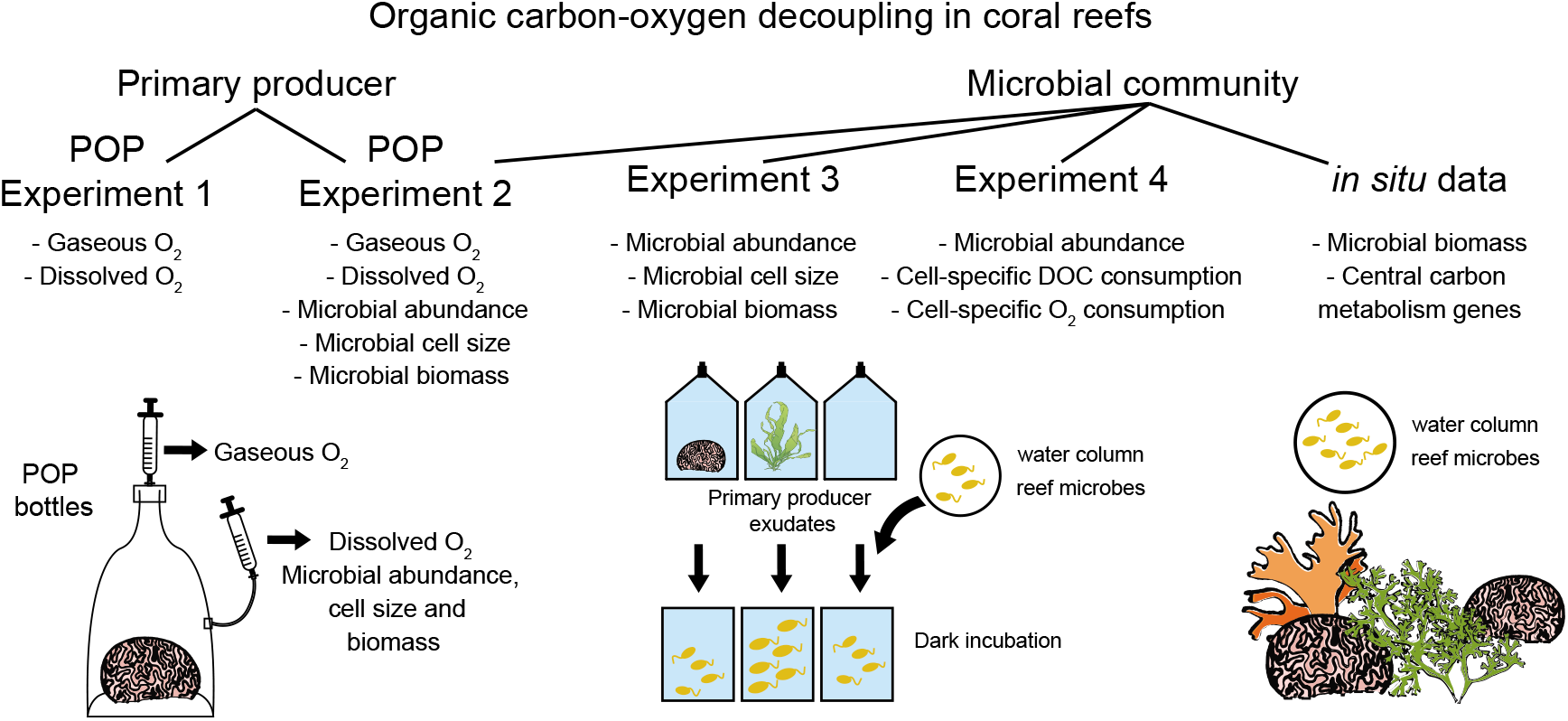
Experimental design for the analysis of metabolic decoupling in coral reef organisms. Peripheral Oxygen Production (POP) Experiments focused on primary producer metabolism, and POP bottles utilized in the experiments are depicted. Microbial growth parameters were analyzed in POP Experiment 2 and dark incubations in Experiments 3 and 4. Microbial biomass and pathways for carbon metabolism were analyzed in reef water samples. The specific variables analyzed in each experiment are listed below experiment number.

POP Experiment 1: The scleractinian coral *Montipora sp*. and the fleshy macroalgae *Chaetomorpha sp*. were collected from coral and macroalgae long-term holding tanks maintained at SDSU. Specimens were collected from the tanks immediately prior to the experiment and placed inside POP bottles. Three specimens of each organism were individually incubated for 4 days with artificial seawater from the coral tank, along with three control bottles under cycles of 12 hours light (125 μmol quanta m^-2^ s^-1^) and 12 hours dark at 27 °C.
POP Experiment 2: Four benthic primary producers were analyzed: the scleractinian coral *Orbicella faveolata*, CCA, the fleshy macroalgae *Chaetomorpha sp*., and turf algae. *O. faveolata* colonies were collected at 12 m depth from the Water Factory site (12°10’91” N, 68°95’49” W) and cut into ~ 10 cm^2^ fragments. Coral fragments were kept for two weeks at the CARMABI Research Station flow-through tank system. The tank was subject to natural diel light cycles with light intensities comparable to 10 m depth, as measured using HOBO Pendant UA-002-64. CCA, turf, and macroalgae were collected off CARMABI immediately prior to the experiment. Five individual incubations for each organism were conducted. Surface area of organisms are provided in Supplementary Dataset 1. Incubations lasted for 2 days at 24 °C with natural diel light cycles.

### Bacterial growth on primary producer exudates

Bacterial growth data, including changes in abundance, cell size and total microbial biomass, derived from three independent experiments: POP Experiment 2, Experiments 3 and 4 (Figure 1). Details on each one of these datasets are given below:

POP Experiment 2: At the start and end of the incubation (time 0h and 48h), 1 mL samples were collected and analyzed by fluorescence microscopy according to McDole et al. 2012. Briefly, cell volume was calculated by considering all cells to be cylinders with hemispherical caps and individual microbial cell volumes were converted to mass in wet and dry weight using previously established size-dependent relationships for marine microbial communities (Simon and Azam 1989). Differences in cell abundance, cell volume, and total microbial biomass were calculated by the difference between final and initial values.
Experiment 3: Five specimens of the coral *Favia sp*. were obtained from a long-term holding tank at the Hawaiian Institute of Marine Biology (HIMB) and placed in independent 5 L polycarbonate containers filled with 0.2 μm-filtered seawater. Five specimens of the fleshy macroalgae *Gracilaria sp*. were collected off HIMB and placed in independent 5 L containers. Additional control containers were filled with filtered seawater. Primary producers were incubated in natural light conditions for 8 hours to release exudates. At the end of the incubation, 2 L of seawater containing exudates were 0.2 μm-filtered and inoculated with 1 L of unfiltered offshore seawater containing water column reef microbial communities. All bottles were incubated for 24 hours in the dark. For microbial abundance and biomass determination, 1 mL samples were collected and analyzed as described above. Differences in cell abundance, cell volume and total heterotrophic microbial biomass were calculated by the difference between final and initial values.
Experiment 4: This dark incubation experiment was performed in 2010 in Mo’orea, French Polynesia. The data was first published in Haas et al. 2011 and was re-analyzed here to investigate cell-specific carbon and O_2_ consumption. Briefly, exudates from corals and algae released during a light incubation period were 0.2 μm-filtered and inoculated with reef microbes. Inoculation in filtered seawater was utilized as control. All bottles were kept in the dark for 48 hours. DOC, dissolved O_2_ and microbial abundances were measured at the beginning and at the end of each dark incubation as described above. DOC samples were filtered through pre-combusted GF/F filters (Whatman, 0.7 μm nominal particle retention) and transferred to acid-washed HDPE vials. Samples were kept frozen until analysis according to Carlson et al. 2010 (Carlson et al. 2010). DOC and O_2_ consumption rates normalized by the bacterial cell yield at the end of each experiment resulted in cell-specific carbon and O_2_ demands. The fleshy organisms used in this experiment were turf algae and the macroalgae *Turbinaria ornata* and *Amansia rhodantha*, and the calcifying organisms were CCA and the coral *Porites lobata*.

### Central carbon metabolism pathways

Microbial metagenomes were sequenced from water samples collected on coral reefs in the Pacific and in the Caribbean. The Pacific samples were collected during NOAA RAMP cruises from 2012-2014 to Hawaiian Islands, Line Islands, American Samoa, and Phoenix Islands. Caribbean samples were collected around the island of Curaçao during the Waitt Institute Blue Halo expedition in 2015. Geographic coordinates for each sampling site are provided in Supplementary Dataset 5, along with benthic cover data. At each sampling site SCUBA divers collected water from within 30 cm of the reef surface using Hatay-Niskin bottles (Haas et al. 2014a). Samples were brought to the ship, filtered through a 0.22 μm cylindrical filter within 4 hours, and stored at -20 °C until laboratory processing. DNA was extracted from the filters using Nucleospin Tissue Extraction kits (Macherey Nagel, Germany) and sequenced on an Illumina HiSeq platform (Illumina, USA). Pacific microbial metagenomes described here were previously analyzed for the presence of CRISPR elements, competence genes and Shannon diversity in Knowles, Silveira *et al.* 2016. Original fastq files from Pacific and Curaçao metagenomes were quality filtered in BBTools with quality score > 20, duplicate removal, minimum length of 60 bases, and entropy 0.7 (Bushnell 2014). The metagenomes are available on the NCBI Short Read Archive (PRJNA494971). Relative abundances of genes encoding rate-limiting enzymes in central carbon metabolism pathways were utilized as proxy for the representation of that pathway in the community. The list of pathways and enzymes analyzed here is provided in Supplementary Table 1. Enzyme-specific databases were built using amino acid sequences from the NCBI bacterial RefSeq. BLASTx searches were performed using metagenomic reads against each enzyme database, using a minimum alignment length of 40, minimum identity of 60 % and e-value < 10^-5^. Reads mapping to the database were normalized by total high-quality reads resulting in the percent abundance of each enzyme. Relative gene abundances are provided as Supplementary Dataset 5. Abundance of bacterial genera in metagenomes were generated through k-mer profiles using FOCUS2 (Silva et al. 2016).

### Statistical tests

All statistical analyses were performed using the software R. The response variables from the incubation experiments were tested for normality using the Shapiro–Wilk test. Due to lack of normality (Shapiro–Wilk, p < 0.05), the nonparametric Kruskal-Wallis test was used to test if there were differences in treatments followed by the post-hoc Wilcoxon test with the False Discovery Rate (FDR) correction with a significance cutoff of p < 0.05. Because gaseous O_2_ production and fraction of total O_2_ in the form of gas were not significantly different between POP Experiments 1 and 2 (Kruskal-Wallis, p > 0.05), we combined the results of both experiments in subsequent tests. For analysis of *in situ* data, microbial biomass was log-transformed to meet the assumption of normality for testing the significance of the slope in the linear regression of fleshy algae cover on microbial biomass. The relationships between fleshy algae cover and enzyme abundances or taxonomic profiles were analyzed by the statistical learning method of supervised random forests. The significance of the random forests variable importance was determined by a permutation method applied to the supervised random forests using the R package rfPermute.

## Results

### One third of algal photosynthetic O_2_ is lost as gas

We designed new incubation chambers called POP (Peripheral Oxygen Production) bottles to quantify the dissolved and gaseous O_2_ release from coral reef primary producers (Figure 1). Dissolved O_2_ production was significantly different among primary producer treatments (Figure 2A, Kruskal-Wallis χ^2^(4) = 19.5, p < 0.05, Supplementary Dataset 1). Primary producers showed higher dissolved O_2_ production compared to controls (Wilcoxon p < 0.05 for all FDR-corrected pairwise comparisons against control). Dissolved O_2_ production was highly variable and not significantly different between fleshy and calcifying organisms (5.99 ± 2.29 and 6.33 ± 2.20 μmol cm^-2^ day^-1^ for fleshy and calcifying, respectively, mean ± SE; Wilcoxon p > 0.05). Gaseous O_2_ production was observed in most primary producer bottles, while no gas was observed in control bottles (Figure 2B, Kruskal-Wallis χ^2^(4) = 30.2, p < 0.05, Wilcoxon p < 0.05 for all pairwise comparisons against control). Fleshy organisms combined produced significantly more gaseous O_2_ than calcifying (0.42 ± 0.35 and 2.33 ± 1.35 μmol cm^-2^ day^-1^ for calcifying and fleshy, respectively, mean ± SE; Wilcoxon p < 0.05). Fleshy macroalgae had the highest fraction of total photosynthetic O_2_ (sum of dissolved and gaseous O_2_) released in the form of gas (37.33 ± 8.34 %), followed by turf algae (13.78 ± 1.33 %), CCA (10.19 ± 2.88 %), and corals (5.00 ± 5.55 %) (Figure 2C). The difference in the fraction of O_2_ as gas was significant among treatments (Kruskal-Wallis χ^2^(4) = 29.2, p < 0.05) and higher in fleshy organisms (Kruskal-Wallis χ^2^(2) = 26.3, p < 0.05; Wilcoxon p < 0.05 for fleshy vs calcifying).

**Figure 2.**
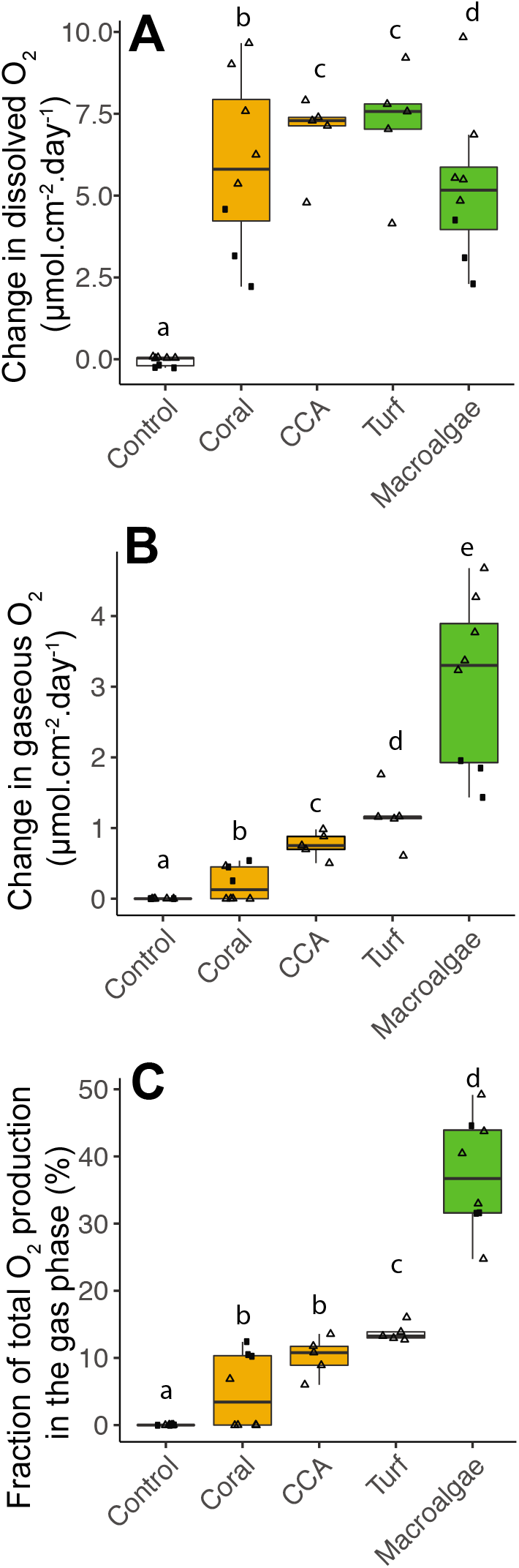
Photosynthetic O_2_ loss as gas from fleshy macroalgae. A) Dissolved O_2_ production rates normalized by organism surface area. B) Gaseous O_2_ production rates normalized by organism surface area. C) Fraction of total photosynthetic O_2_ production in the form of gas. Solid squares correspond to Experiment 1 and triangles to Experiment 2. Primary producer had a significant effect on all three variables (Kruskal-Wallis p < 0.05), and letters above boxes indicate p < 0.05 for Wilcoxon pairwise tests with FDR correction. Orange indicate calcifying and green fleshy organisms.

### Bacteria accumulate more carbon under high DOC:O_2_

During POP Experiments, the change in bacterial abundance in primary producer bottles was higher compared to controls (Figure 3A, Kruskal-Wallis χ^2^(4) = 20.6, p < 0.05, Wilcoxon p < 0.05 for all pairwise comparisons against control, Supplementary Dataset 2). The change in abundance was higher in fleshy algae treatments compared to calcifying treatments (Kruskal-Wallis χ^2^(2) = 16.4, p < 0.05, Wilcoxon p < 0.05 for pairwise comparison of fleshy vs calcifying). Microbial cell volume changed differently in treatments during incubations (Figure 3B, Kruskal-Wallis x^2^(5) = 1235.2, p < 0.05, Supplementary Dataset 3). Bacteria growing in control bottles showed no change in volume (Wilcoxon, p > 0.05 for pairwise comparison of control vs T0h). When growing on coral and CCA exudates, bacteria became smaller, while growing on turf and macroalgae exudates lead to an increase in bacterial cell volume (Wilcoxon p < 0.05 for all pairwise comparisons vs T0h and vs control). When taking abundance and cell volume into account, the change in total bacterial biomass was significantly different among treatments (Figure 3C, Kruskal-Wallis χ^2^(4) = 21.6, p < 0.05, Supplementary Dataset 2). Coral exudates decreased total bacterial biomass, while turf and macroalgae increased total bacterial biomass, and CCA lead to no change (Wilcoxon, p < 0.05 for pairwise comparisons of coral, turf and macroalgae vs control). The change in biomass in the two fleshy algae together was greater than in the calcifying organisms together (Kruskal-Wallis χ^2^(2) = 16.6, p < 0.05; Wilcoxon, p < 0.05 for pairwise comparison of fleshy vs calcifying).

**Figure 3.**
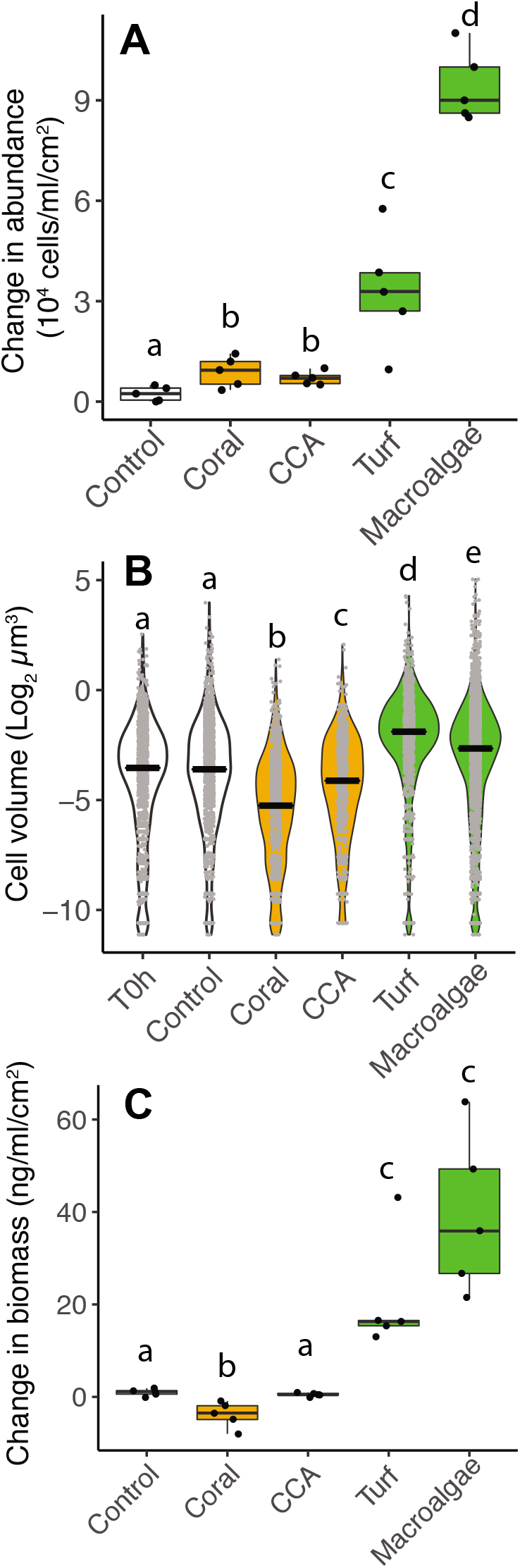
Differences in microbial biomass accumulation in primary producer exudates. A) Changes in microbial abundance normalized by primary producer surface area. B) Cell volume distributions. C) Changes in total microbial biomass, accounting for abundance and cell volume, normalized by primary producer surface area. Primary producer treatments had a significant effect on all three microbial variables (Kruskal-Wallis p < 0.05), and letters above boxes indicate p < 0.05 for Wilcoxon pairwise tests with FDR correction. Orange indicate calcifying and green fleshy organisms.

A bottle incubation performed at the Hawaiian Institute of Marine Biology (HIMB) with a distinct set of primary producers showed the same pattern in microbial physiology (Experiment 3, Supplementary Datasets 2 and 3). After 24 hours of dark incubation on exudates but in the absence of primary producer, microbial abundances increased in both the coral and algae treatments (Figure S1A, Kruskal-Wallis χ^2^(2) = 7.2, p < 0.05). Changes in cell volume were not significant among treatments (Figure S1B, Kruskal-Wallis χ^2^(2) = 3.8, p > 0.05). However, the resulting change in total microbial biomass when accounting for cell volume and abundance showed an increase in biomass in macroalgae treatments, while coral treatments did not change and controls decreased in biomass (Kruskal-Wallis χ^2^(2) = 7.2, p < 0.05).

### Bacteria decouple DOC and O_2_ consumption

Cell-specific carbon and O_2_ consumption by heterotrophic metabolism was obtained by normalizing the change in DOC and O_2_ concentrations by the cell yield in a dark incubation (Experiment 4 in Figure 1, Supplementary Dataset 4). The amount of DOC consumed per cell was higher in fleshy macroalgae treatments compared to coral and control treatments (i.e., low cell yields per carbon consumed, Figure 4A, Kruskal-Wallis χ^2^(2) = 13.7, p < 0.05; Wilcoxon, p < 0.05 for pairwise comparisons of macroalgae vs controls and vs corals). The amount of O_2_ consumed per cell was not different between treatments (Figure 4B, Kruskal-Wallis χ^2^(2) = 0.5, p > 0.05).

**Figure 4.**
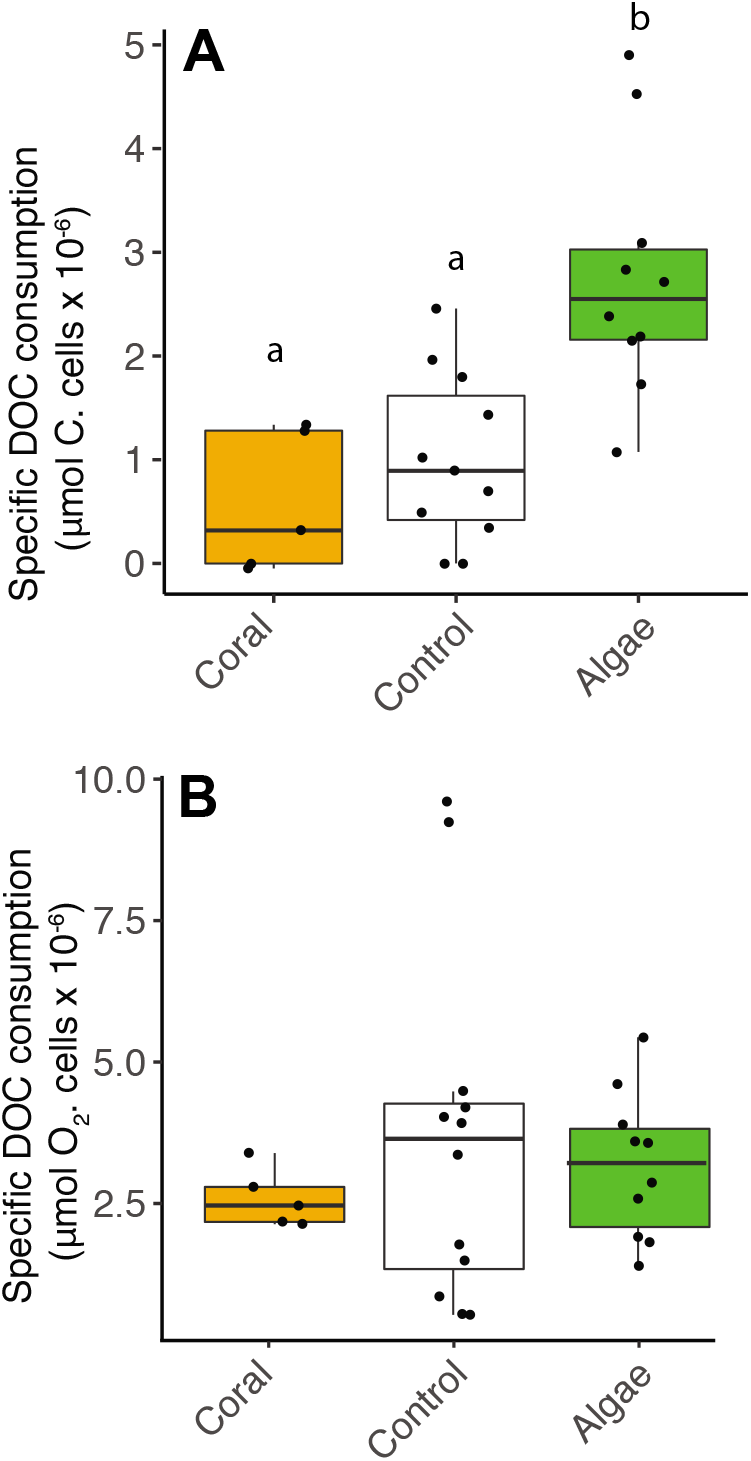
Decoupling between microbial DOC and O_2_ consumption in fleshy macroalgae exudates. Cell-specific carbon and O_2_ consumption data from Experiment 4: dark incubations of microbial communities in primary producer exudates. A) Cell-specific DOC consumption. B) Cell-specific O_2_ consumption. Primary producer treatments had a significant effect on specific DOC consumption only (Kruskal-Wallis p < 0.05) and letters above boxes indicate p < 0.05 for Wilcoxon pairwise tests with FDR correction. Orange indicate calcifying and green fleshy organisms.

### Warburg-like metabolism causes bacterial biomass accumulation

The relative abundance of genes encoding rate-limiting enzymes of central carbon metabolism pathways was quantified in microbial metagenomes from reefs across a gradient in algae cover (Supplementary Dataset 5). In this dataset, water column microbial biomass increased with fleshy algae cover (Linear regression, p < 0.05, m = 0.009, R^2^ = 0.29 for the comparison between log10-transformed biomass and fleshy algae cover). The random forest analysis with genes encoding rate-limiting enzymes of central carbon metabolism explained 14.3 % of the fleshy algae cover. The genes with highest explanatory power were phosphoenolpyruvate carboxylase, aspartate aminotransferase, oxoglutarate dehydrogenase, glucose 6-phosphate dehydrogenase and KDPG aldolase (increasing mean squared error 15.78, 9.73, 9.21, 8.70 and 6.95 %, respectively). The relationships between these enzymes and fleshy algae cover are shown in Figure 5.

**Figure 5.**
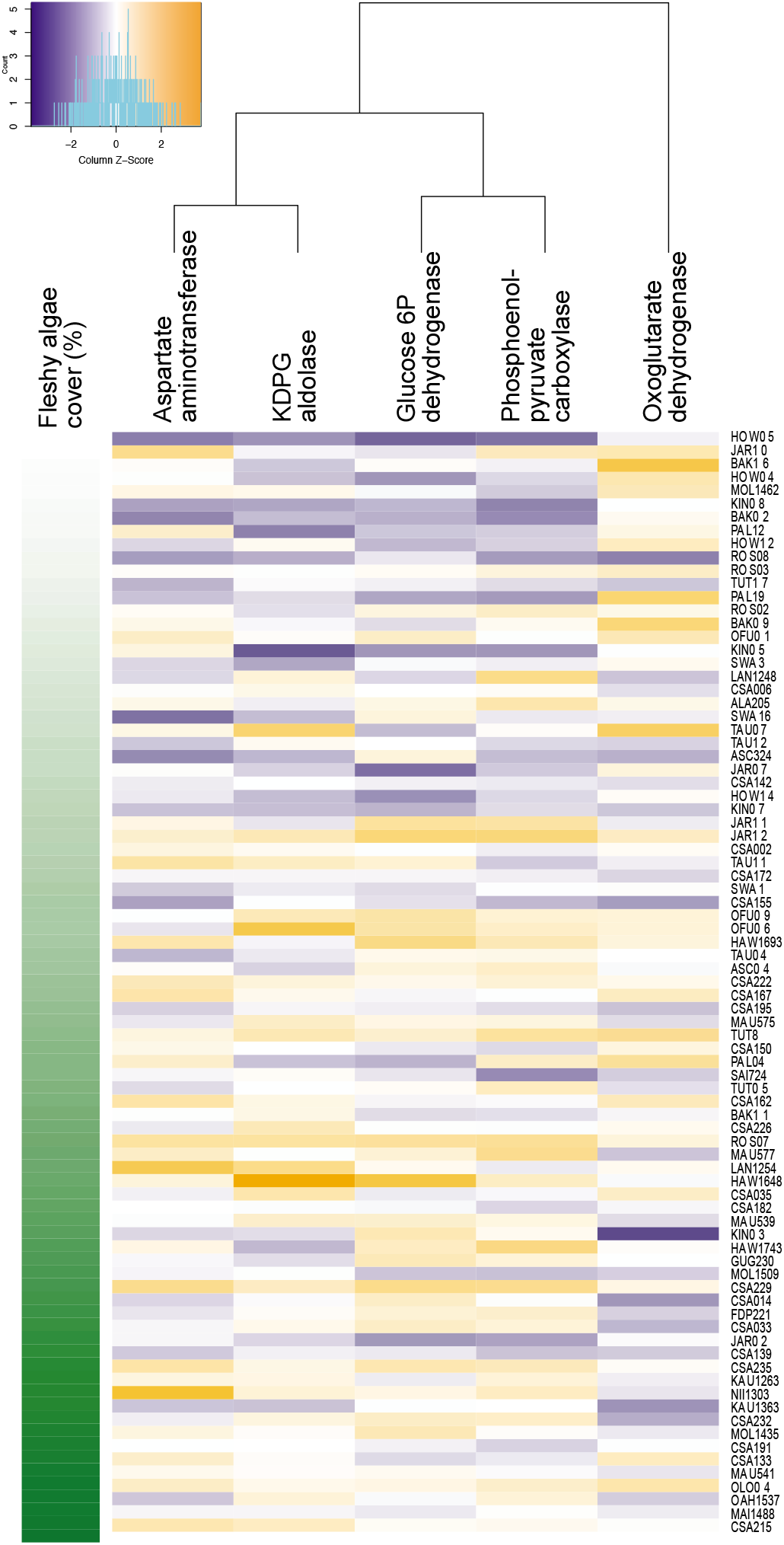
Increase in genes encoding anabolic pathways and decrease in genes encoding carbon oxidation reactions with fleshy algae cover *in situ*. Relative abundances of genes encoding rate-limiting enzymes in reef metagenomes are plotted in relation to fleshy algae cover (sum of fleshy turf and macroalgae). Individual metagenomes are listed as rows and sorted by the fleshy algae cover, ranging from 15 to 94.1 %. Enzyme abundance was scaled by column to allow between-enzyme comparisons. Only enzymes significantly predicting fleshy algae cover in the random forest analysis are shown. A complete list of enzymes and abundances is provided as Supplementary Data 5.

Phosphoenolpyruvate carboxylase and aspartate aminotransferase are involved in anaplerotic reactions, glucose 6-phosphate dehydrogenase diverts glucose to both Entner-Doudoroff and Pentose Phosphate pathways, and KDPG aldolase is unique to the Entner-Doudoroff pathway. These four enzymes had a positive relationship with fleshy algae cover (random forest mean decreasing accuracy, p < 0.05). Oxoglutarate dehydrogenase is an oxidative enzyme in the Krebs cycle, and its relative abundance decreased with fleshy algae cover (random forest mean decreasing accuracy, p < 0.05).

To test if these changes in relative gene abundances could be explained by the taxonomic dominance of bacterial genera encoding these genes, we tested the relationship between fleshy algae cover and taxonomic profiles at the genus level using random forest. While several taxonomic groups changed their abundance with increasing fleshy algae cover, none of these groups consistently lack or encode the enzymes of the central carbon pathways described here, with high strain-level variation in their functional genetic repertoire (Supplementary Dataset 5 shows the relative abundances of genera with significant relationship with fleshy algae cover in the random forest mean decreasing accuracy, p < 0.05).

## Discussion

### O_2_ ebullition modifies reef biogeochemistry

Photosynthesis and respiration have a theoretical 1:1 molar ratio of carbon and O_2_ produced and consumed (Williams et al. 1979). In holobionts with a large heterotrophic component, such as corals, respiration consumes at least 50 % of the oxygen and organic carbon produced in photosynthesis (Tremblay et al. 2016). Respiration quotients for symbiotic cnidarians range from 0.8 and 0.9, consuming more O_2_ than carbon (Muscatine et al. 1981). Based on these processes, the higher ratios of DOC:O_2_ observed in fleshy algal exudates compared to corals are counter-intuitive (Haas et al. 2011). O_2_ ebullition is a plausible explanation for this observation. The organic carbon released by algae stays in solution, while a fraction of the O_2_ nucleates and escapes, leaving behind a high DOC:dissolved O_2_ ratio. Differential carbon allocation into biomass can also alter these ratios, and the quantitative analysis of carbon incorporation in combination with dissolved and gaseous O_2_ dynamics is the next step to resolve these budgets.

O_2_ ebullition and bubble injection by hydrodynamics are recognized as a source of error when estimating production in shallow water ecosystems, yet the extent to which ebullition affects these estimates is rarely quantified due to methodological challenges (Kraines et al. 1996; Cheng et al. 2014). O_2_ ebullition observed in this study corresponded to 5 – 37 % of net primary production and accounted for the loss of up to 21 and 37 % of photosynthetically produced O_2_ in a salt marsh and a lake, respectively (Koschorreck et al. 2017; Howard et al. 2018). If *in situ* bubble nucleation and rise rates are comparable to those in incubations, our results imply that coral reef primary production has been significantly underestimated (Howard et al. 2018). In coral reefs, bubbles were reported on the surface of turf algae, on sediments, and inside coral skeletons colonized by endolithic algae (Odum and Odum 1955; Bellamy and Risk 1982; Clavier et al. 2008; Freeman et al. 2018). The heterogeneous distribution of bubble nucleation over primary producers entail that benthic community compositional shifts determine the magnitude of O_2_ ebullition on reef-level O_2_ dynamics. A combination of *in situ* incubations and gas exchange studies is required to establish such relationships.

### Warburg effect is the biophysical mechanism of microbialization

High DOC:O_2_ release ratio by algae reflect bacterial carbon and oxygen metabolism. When growing on algal exudates, microbes increased cell volume and DOC consumption per cell (Figure 3 and 4A). These cells displayed the same cell-specific O_2_ consumption compared to small cells growing on coral exudates (Figure 4B). These results indicated that microbes growing on coral exudates fully oxidize the organic carbon consumed, channeling metabolic energy to maintenance costs (Russell and Cook 1995). On algal exudates, bacteria had higher cell-specific DOC consumption, with greater fraction of the organic carbon shunted to biosynthesis (incomplete oxidation), and less relative O_2_ consumption, culminating in increased community biomass. This process is classically described as the Warburg effect in eukaryotic cells undergoing fast metabolism and limited by the enzyme catalytic rates, as opposed to substrate concentrations (Vander Heiden et al. 2009). In bacteria, the Warburg effect is a physiological response that optimizes energy biogenesis and biomass synthesis in face of changing proteomic demands (Basan et al. 2015). This biochemical transition occurs because the proteome cost of energy biogenesis by respiration exceeds that by fermentation.

The decoupling between bacterial specific carbon and O_2_ consumption observed in Figure 4 predicts an increase in pathways that shunt carbon to biosynthesis and incomplete oxidation when bacteria are growing on algal exudates (Russell and Cook 1995; Haas et al. 2016). The Pentose Phosphate (PP) and Entner-Doudoroff (ED) pathways are alternatives to the canonical glycolytic route for carbohydrate consumption (Embden-Meyerhoff-Parnas, EMP). Both PP and ED produce more NADPH than NADH. These two reduced coenzymes have distinct functions: NADPH is preferentially consumed in biosynthetic routes for lipid, amino acid, and nucleotide synthesis while NADH is preferentially consumed in oxidative phosphorylation for ATP production (Pollak et al. 2007; Spaans et al. 2015). Therefore, cells utilizing more NADH pathways will consume more O_2_ relative to carbon compared to cells utilizing more NADPH. Our metagenomic data is in line with this prediction. The abundance of genes encoding glucose-6-phosphate dehydrogenase and KDPG aldolase increased with fleshy algae cover (Figure 5). Glucose-6-phosphate dehydrogenase deviates glucose to both the PP and ED pathways, and KDPG aldolase is unique to ED.

The tricarboxylic acid cycle (TCA or Krebs cycle), is a canonically oxidative path for complete decarboxylation of pyruvate. But the TCA cycle can work as a biosynthetic pathway by providing intermediates to amino acids, nitrogenous bases and fatty acids syntheses (Sauer and Eikmanns 2005; Cronan and Laporte 2013). TCA cycle stoichiometry is maintained through the resupply of these intermediates by anaplerotic reactions that use pyruvate (or phosphoenolpyruvate) (Owen et al. 2002). Here we found an increase in the abundance of genes encoding anaplerotic enzymes and decrease in genes encoding oxoglutarate dehydrogenase, catalyzes an oxidative decarboxylation in the Krebs cycle, with increasing algal cover. These results indicate a shift towards the use of Krebs cycle as an anabolic route (Sauer and Eikmanns 2005; Bott 2007). Anaplerotic activity increases in rapidly growing bacteria with high rates of amino-acid synthesis (Bott 2007). These routes shunt organic carbon into biomass accumulation, as opposed to oxidation with electron transfer to O_2_ (Russell and Cook 1995). This is consistent with the bacterial growth and DOC consumption patterns observed in our incubation experiments.

Alternative hypotheses for DOC and O_2_ consumption patterns are *broken* TCA cycles and reactive oxygen species (ROS) detoxification (Mailloux et al. 2007; Steinhauser et al. 2012). There was no evidence for an increase in genes encoding these pathways in our dataset (Supplementary Data 5). The metagenomic transitions observed cannot be explained by the rise or disappearance of specific taxonomic groups associated with microbialization, such as *Vibrio, Flavobacteria* and SAR11 (Dinsdale et al. 2008; Kelly et al. 2014). The genes encoding the pathways discussed here are not consistently encoded by these taxonomic groups, and the functional genomic transitions observed here are likely a result of strain-level compositional shifts not directly related to the taxonomic structure of the community at genus or family level, as is often the case for coral reefs (Klingner et al. 2015; Silveira et al. 2017b).

### O_2_ loss affects ecosystem biomass allocation

O_2_ depletion observed in previous dark incubations with algae exudates was a result of high bacterial densities, and not of high cell-specific respiration rates (Haas et al. 2016). The cell volume and abundance changes observed in our experiments explain *in situ* microbial biomass (McDole et al. 2012; Haas et al. 2016). In degraded reefs, fleshy algae release more exudates than corals, fundamentally modifying the biogeochemical environment for heterotrophic metabolism (Figure 6). The ensuing increases in bacterial size, abundance and DOC consumption described in our incubations make bacterial energetic demands surpass that of macrobes, and explain the observed decrease in DOC standing stocks in degraded reefs (McDole et al. 2012; Haas et al. 2016; Somera et al. 2016). Increase in overflow metabolism due to the decoupling between anabolic and catabolic reactions is consistent with the high amounts of particulate material observed in algae-dominated reefs, affecting reef-scale trophic interactions (Russell and Cook 1995; Wilson et al. 2003; De’ath and Fabricius 2010; Silveira et al. 2015; Haas et al. 2016). The metabolic decoupling described here is also likely to occur in planktonic oceanic systems in response to global changes: warmer, high pCO_2_ waters depleted of nutrients due to stratification shift primary producer community composition, increasing the release of carbon-rich exudates by phytoplankton while decreasing O_2_ (Boyd et al. 2010; Oschlies et al. 2018). Pelagic heterotrophic bacterial metabolism display a positive relationship between substrate energy density and turnover times, with no relationship with respiration rates (Casey et al. 2015). This pattern is consistent with metabolic shifts described here, decreasing microbial biomass turnover rates in the oceans (Vallino et al. 1996).

**Figure 6.**
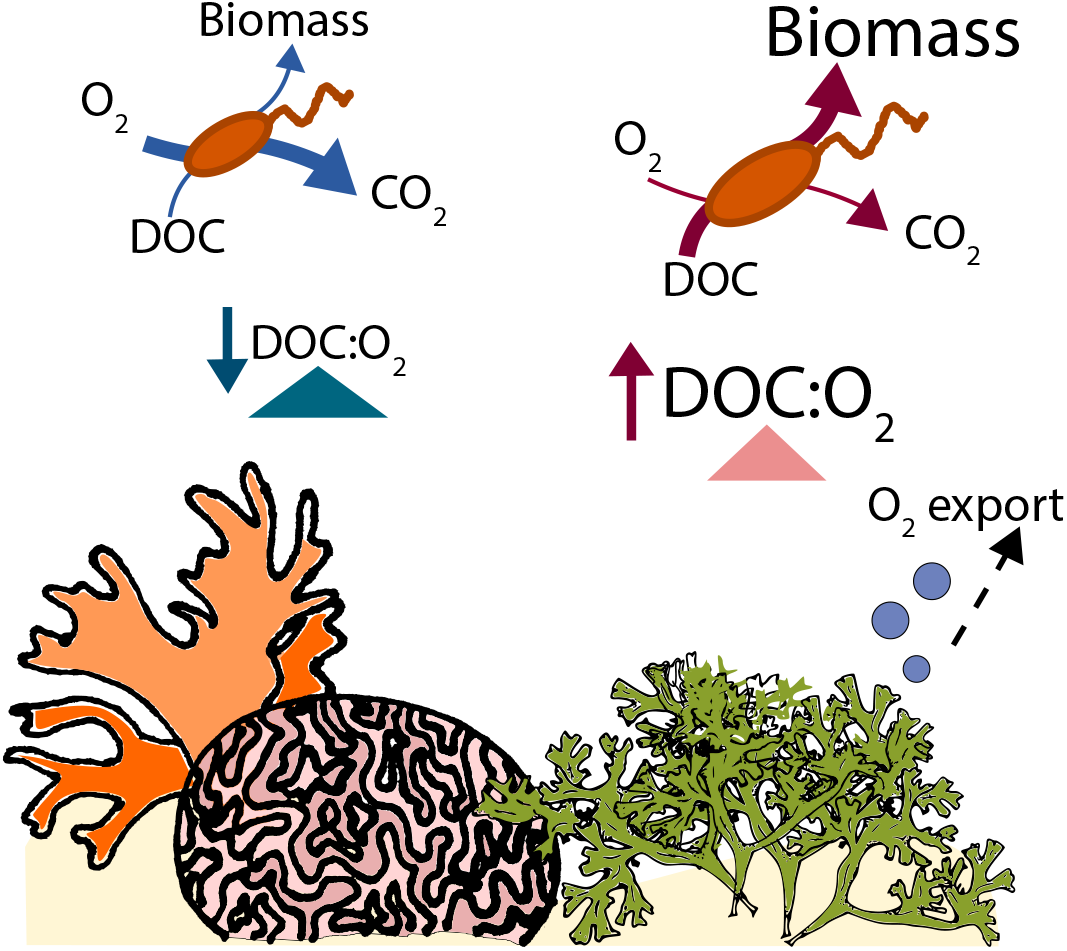
Metabolic decoupling model. Fleshy algal exudates have higher DOC:O_2_ ratios compared to corals. Heterotrophic bacteria respond to this difference by decoupling organic carbon consumption from oxidation: higher DOC consumption per cell with no increase in O_2_ consumption per cell. This metabolic change leads to accumulation of microbial biomass.

## Conclusion

Ebullition causes a decoupling between photosynthetic fixed carbon and O_2_ in fleshy macroalgae. The O_2_ loss by ebullition fundamentally changes the reef biogeochemical environment. In contrast, there is almost no O_2_ loss through ebullition associated with corals, because the zooxanthellae reside in the tissue of the animal. On reefs with large amounts of fleshy macroalgae, the microbial community is dominated by anabolic metabolisms, with corresponding increases in cell volume, abundance and specific DOC consumption pathways. These physiological changes cause microbial biomass accumulation with detrimental effects on coral health and trophic relationships.

## Supporting information

## Acknowledgements

We thank the crew and captain of the NOAA ship Hi’ialakai and Waitt Institute ship Plan B that contributed to sampling. We thank the CARMABI and HIMB teams for the support conducting field experiments. We thank Robert Edwards at SDSU for access to computer servers funded by the NSF (CNS-1305112 to Robert A. Edwards). This work was funded by the Gordon and Betty Moore Foundation (grant 3781 to FR), Spruance Foundation, NOAA, and the Waitt Institute. CBS was funded by CNPq (234702 to CBS) and Spruance Foundation. TNFR was supported by the NSF (G00009988 to TNFR).

## Conflict of interest statement

The authors declare no conflict of interest.

