## Supplementary material for "Biophysical and physiological causes of coral reef microbialization"

Silveira et al. 2018

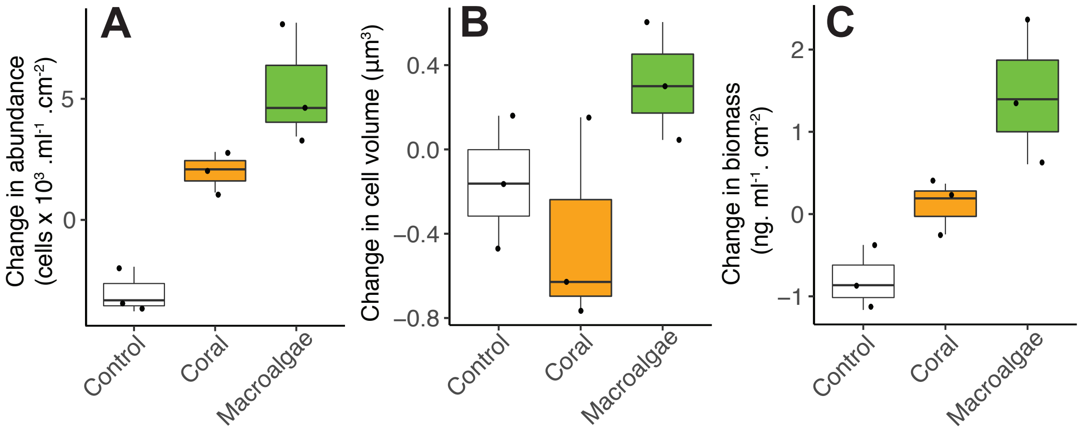


**Figure S1.** Changes in microbial abundance (A), cell volume (B) and total biomass (C) stimulated by coral and algal exudates in Experiment 3.

**Table S1.** List of central enzymes in central carbon metabolism, reactive oxygen species detoxification and replication/translation analyzed in this study. Gene abundances are provided in Supplementary Data 5.

| Pathway | Rate-limiting enzymes | Enzyme Commission (EC) number |
|---|---|---|
| Anaplerosis | Pyruvate carboxylase | 6.4.1.1 |
| | Phosphoenolpyruvate carboxylase | 4.1.1.31 |
| | Phosphoenolpyruvate carboxykinase | 4.1.1.49 and 4.1.1.32 |
| | Glutamate dehydrogenase | 1.4.1.2 and 1.4.1.4 |
| | Aspartate aminotransferase | 2.6.1.1 |
| Embden–Meyerhof–Parnas | Phosphofructokinase | 2.7.1.11 |
| | Pyruvate kinase | 2.7.1.1 |
| Entner-Doudoroff | 2-keto-3-deoxygluconate-6-phosphate (KGDP) aldolase | 4.1.2.14 |
| Pentose Phosphate | 6-phosphogluconate dehydrogenase | 1.1.1.44 |
| Reactions fueling Entner-Doudoroff and Pentose Phosphate | Glucose 6P dehydrogenase | 1.1.1.49 |
| | Glucose dehydrogenase | 1.1.1.47 |
| | Gluconate kinase | 2.7.1.12 |
| Krebs Cycle (oxidative) | Pyruvate dehydrogenase complex (E1) | 1.2.4.1 |
| | Oxoglutarate dehydrogenase | 1.2.4.2 |
| | Isocitrate dehydrogenase | 1.1.1.41 and 1.1.1.42 |
| Glyoxylate cycle | Malate synthase | 2.3.3.9 |
| | Isocitrate lyase | 4.1.3.1 |
| ROS detoxification | Superoxide dismutase | 1.15.1.1 |
| | Glutathione reductase | 1.8.1.7 |
| Transcription | rpoB | RNA polymerase |
